# Classification of columnar and salt-and-pepper organization in mammalian visual cortex

**DOI:** 10.1101/698043

**Authors:** Jaeson Jang, Min Song, Se-Bum Paik

**Author notes:** These authors contributed equally to this work (Jaeson Jang, Min Song). Correspondence to Se-Bum Paik.

## Abstract

In mammalian visual cortex, neural tuning to stimulus orientation is organized in either columnar^1^ or salt-and-pepper^2^ patterns across species. This is often considered to reflect disparate mechanisms of cortical development across mammalian taxa. However, it is unknown whether different cortical architectures are generated by species-specific mechanisms^3,4^, or simply originate from the variation of biological parameters within a universal principle of development^5–8^. We analysed neural parameters in eight mammalian species and found that cortical organization is predictable by a single factor: the retino-cortical mapping ratio. We show that a Nyquist sampling model explains parametric division of the patterns with high accuracy and that simulations of controlled mapping conditions reproduce both types of organization. Our results explain the origin of distinct cortical circuits under a universal development process.

---

Neural tuning to visual stimulus orientation is one of the hallmarks of the primary visual cortex (V1) in mammals. Intriguingly, this tuning in V1 is organized into distinct topographic patterns across species, such as columnar orientation maps in primates^1^ and salt-and-pepper type organization in rodents^2^ (Fig. 1a). From the fact that species of distinct cortical organization are found on separate branches of the mammalian phylogenetic tree, it has been suggested that columnar or salt-and-pepper organization reflect species-specific principles of evolution underlying the development of cortical circuits^3,4^.

**Figure 1.**
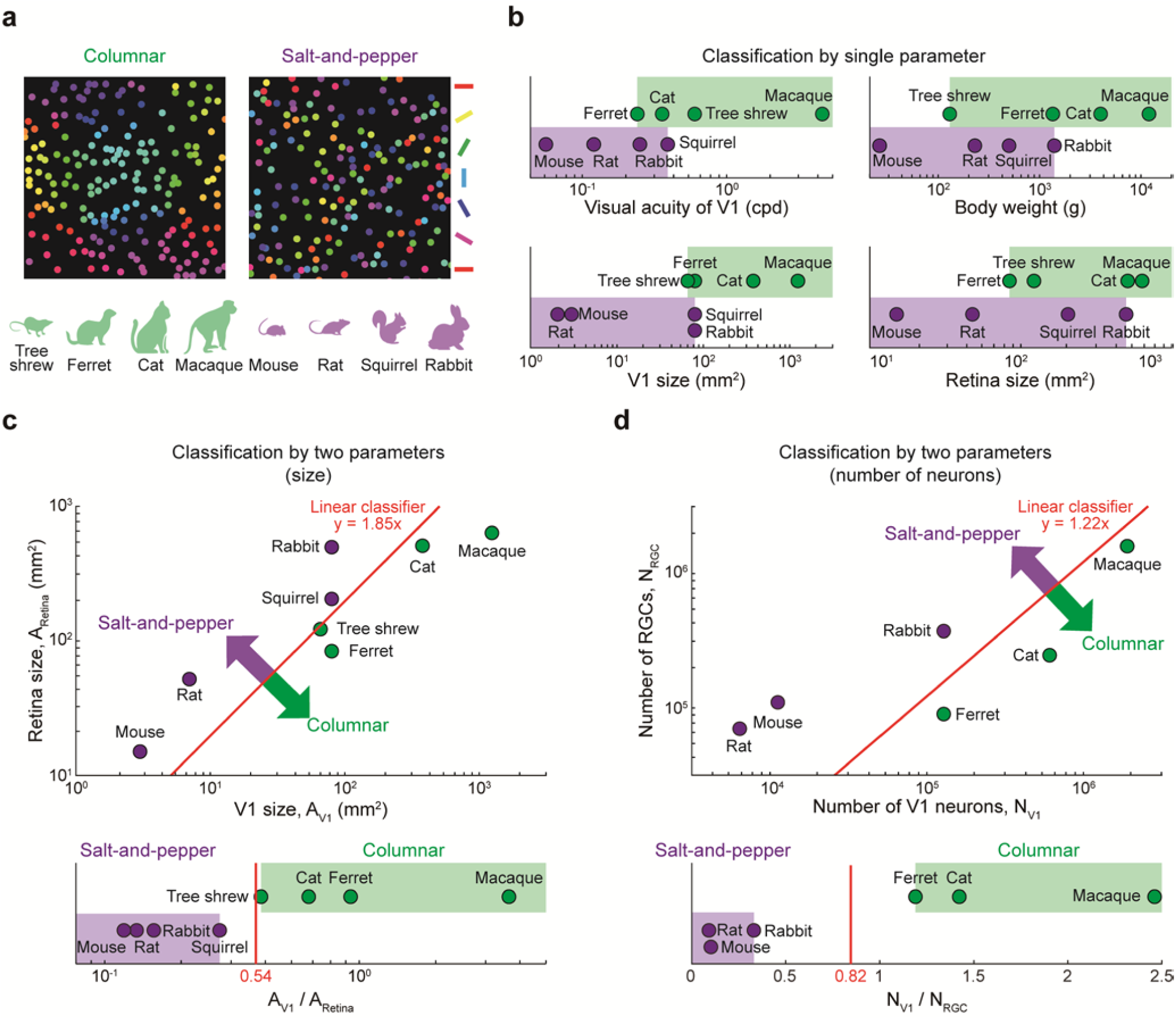
Parametric division of spatial organization of orientation tuning across species. **a**, Columnar and salt-and-pepper organization of orientation tuning observed in mammalian species. **b**, Whether a species has columnar or salt-and-pepper organization cannot be clearly predicted by visual acuity, body weight, V1 size or retina size. **c**, *Top*, V1 organization of eight mammalian species can be divided by a linear classifier (*y* = *ax*) in 2D space as a function of V1 and retina sizes. *Bottom*, V1 organization of each species is well predicted by the ratio between the size of V1 and of a retina. **d**, Similar analysis as in (c) with the number of RGCs and V1 neurons as determinant parameters.

Another recent view is that the cortical development is governed by a universal mechanism, but that disparate architectures can arise from variation of specific biological parameters, such as the range of cortical interaction^5^, the size of V1^6^ or the number of V1 neurons^7^. However, further analysis of data for various species resulted in reported counterexamples of this simple prediction, implying that V1 organization may not be simply determined by a single anatomical factor^8^. For instance, four species of mammals (ferret, tree shrew, rabbit and gray squirrel) have V1 of comparable size, but two of them (ferret and tree shrew) have columnar orientation maps, while the others (rabbit and gray squirrel) have salt-and-pepper organization (Fig. 1b, left bottom). Similarly, other candidate parameters such as visual acuity or body weight also failed to predict V1 organization of orientation tuning among these four species^8^ (Fig. 1b).

Herein, from the analysis of data for eight mammalian species, we propose that the mapping ratio between retina and cortex solely predicts cortical organization across species. We show that distinct cortical circuits may originate from the variation of specific biological parameters under a universal development process and that a Nyquist sampling model explains this parametric division of cortical development with high accuracy.

## Results

We first found that the V1 organization of the eight species reported so far can be successfully divided into columnar maps or salt-and-pepper organization in linear classification (Fig. 1c, top), by considering the size of retina (A_Retina_) and the size of V1 (A_V1_) together (Extended Data Table 1). This analysis showed that the ratio between the size of V1 and of a retina can solely predict the V1 organization in all the test data (Fig. 1c, bottom). In particular, cortical organization of the four species with V1 of similar size (∼80 mm^2^ for ferrets, tree shrews, rabbits and gray squirrels) could also be predicted as columnar or salt-and-pepper organization, which could not be determined by any single biological parameter in previous studies. Then, to examine the mapping between two areas in terms of the neural sampling ratio, we further tried a prediction based on the ratio between the number of RGC (N_RGC_) and V1 neurons (N_V1_) (i.e. retino-cortical sampling ratio) and found that this ratio also successfully predicts V1 organization of all these species (Fig. 1d).

Then, what is the underlying principle that explains this classification of V1 organization by retino-cortical sampling? Based on previous experimental observations that the orientation tuning in V1 can be predicted by the arrangement of ON and OFF feedforward afferents^9,10^, we assumed that both columnar maps and salt-and-pepper organization can arise from the same retinal mosaics, simply from different feedforward mapping ratios (Fig. 2a). Using observed RGC mosaics data in cats^11^, we performed a simulation of cortical organization of orientation tuning using our previous developmental model of orientation maps^12,13^. As a result, we found that both columnar and salt-and-pepper organizations of orientation tuning can develop, even from the same RGC mosaics, only if the sampling ratio between RGC and V1 neurons (N_V1_/N_RGC_) varies (Fig. 2a, top vs. bottom). We simulated two scenarios with two different sizes of the model V1, but with same neuron density. It was assumed that the entire retinal area is matched to the whole V1 patch and that each V1 neuron receives feedforward inputs from the same size of local ON and OFF RGCs in the area of the corresponding retinal location (See Methods for details). We assumed that every cortical cell receives input from a similar number for RGCs regardless of variation in RGCs density and retino-cortical magnification, because the density of the RGCs and retino-cortical magnification are proportional across retinal eccentricity in the observed RGC data at large scale^32,33^ (Extended Data Fig. 1). Then, the orientation tuning of each V1 neuron was calculated from the receptive field of sampled RGCs^14^ (Extended Data Fig. 2).

**Figure 2.**
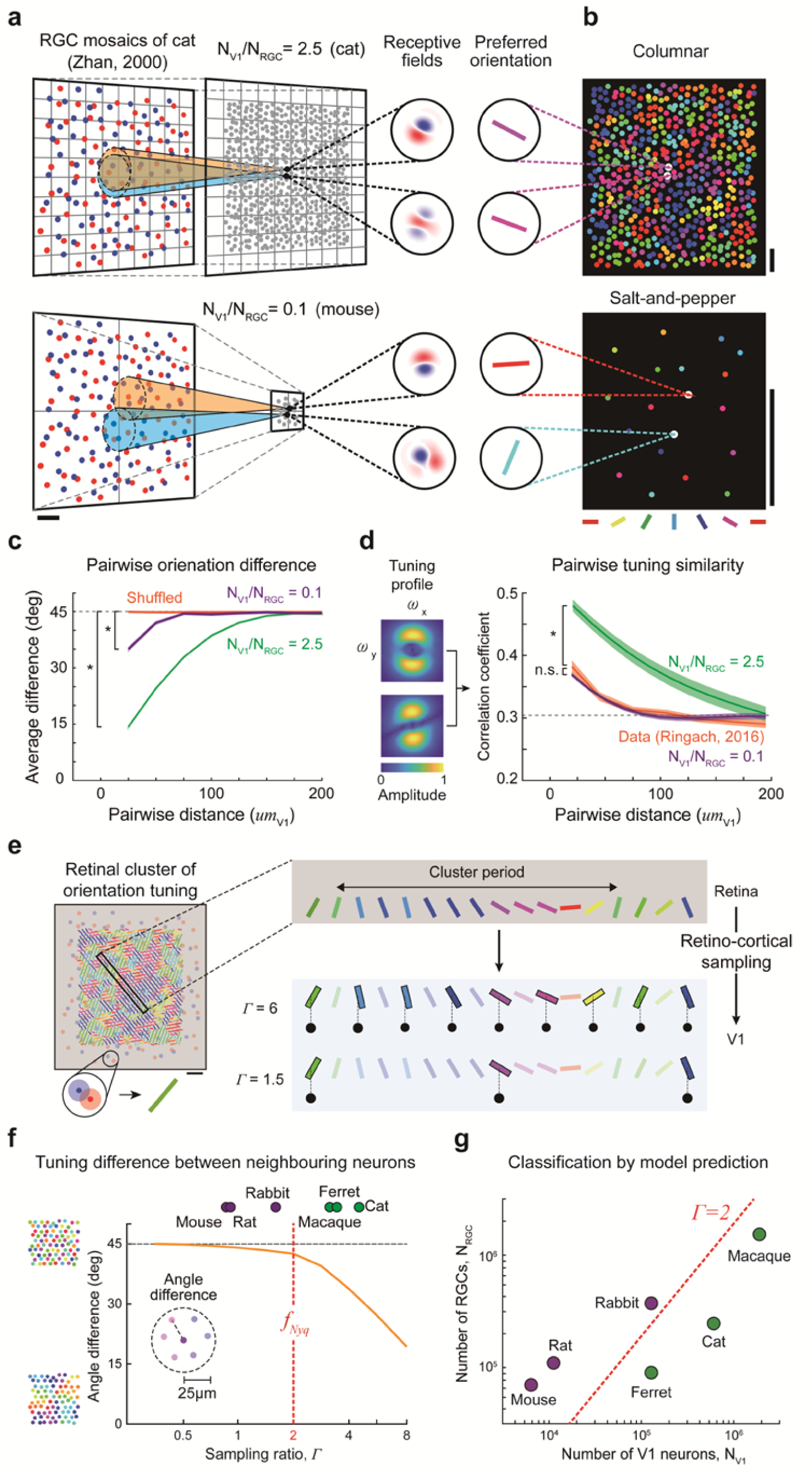
V1 organization as a Nyquist sampling in the retino-cortical projection. **a**, *Top*, For relatively large V1 size, retinotopic mapping has a high sampling density from retina to V1 space, thus neighbouring V1 neurons have highly overlapping receptive fields, resulting in similar orientation preference. *Bottom*, For relatively small V1, neighbouring V1 neurons receive inputs from weakly overlapping RGC populations due to the sparse sampling density, resulting in dissimilar orientation preference. **b**, From the same RGC mosaics profile, clustered map or salt-and-pepper organization is seeded according to the relative size of V1. **c**, Difference between the preferred orientation of neighbouring V1 neurons. N_V1_/N_RGC_ = 0.1 vs. shuffled (violet vs. orange), *p < 0.01; N_V1_/N_RGC_ = 2.5 vs. shuffled (green vs. orange), *p < 0.01; two-sample Kolmogorov-Smirnov test. **d**, Tuning similarity among neighbouring neurons for different mapping conditions. *Left*, Example of tuning profiles of V1 receptive field obtained using FFT analysis. *Right*, Weakly clustered orientation tuning observed in mice^17^ is reproduced by the model (N_V1_/N_RGC_=0.1). Data vs. N_V1_/N_RGC_ = 0.1 (violet and orange), p = 0.29; data vs. N_V1_/N_RGC_ = 2.5 (green and orange), *p < 0.01; two-sample Kolmogorov-Smirnov test **e**, *Left*, Expected cluster pattern of preferred orientation across retinal mosaics in (a). *Right*, According to the Nyquist theorem, high *Γ* (n_V1_ in a single period in retinal mosaics) of cortical sampling induces a continuous change of orientation tuning, while low *Γ* (< 2) would cause an aliasing of RGC periodicity and an abrupt change of orientation tuning. **f**, The model predicts that the organization of cortical orientation tuning makes a sharp transition around the Nyquist sampling frequency (*f*_*Nyq*_), matching the observations of estimated sampling ratios across species. **g**, The groups of species with or without columnar clustering were successfully distinguished by the sampling ratios predicted by the Nyquist theorem. Scale bar, the average distance of OFF RGCs (*d*_*OFF*_) for (a), (e) and 100 µm in cortical space for (b).

When V1 size was relatively large (N_V1_/N_RGC_ = 2.5), neighbouring V1 neurons had highly-overlapping receptive fields due to a high sampling density of mapping from retina to V1 space (Fig. 2a, top). As a result, neighbouring V1 cells had similar orientation tuning (Fig. 2b, top and 2c, green curve). In contrast, when V1 size was relatively small (N_V1_/N_RGC_ = 0.1), neighbouring V1 neurons received inputs from weakly-overlapping RGC populations due to the sparse sampling density of the mapping (Fig. 2a, bottom), thus inducing a salt-and-pepper organization (Fig. 2b, bottom and 2c, violet curve). These results suggest that distinct V1 organization can originate from the same RGC mosaics, but with different retino-cortical sampling ratio according to the size of the retina and V1.

Interestingly, we found that even for low N_V1_/N_RGC_, V1 organization of orientation tuning can be slightly clustered as in mice16,17 (Fig. 2c, N_V1_/N_RGC_ = 0.1 vs. shuffled (violet vs. orange), *p < 0.01; two-sample Kolmogorov-Smirnov test). This is because the sampling of neighbouring V1 neurons can partially overlap in the retinal space^15^. We found a condition of salt-and-pepper organization with weak clustering that quantitatively matched the statistics of pairwise tuning similarity in recent observations of salt-and-pepper organization in mice^16,17^ (Fig. 2d, data vs. N_V1_/N_RGC_ = 0.1 (violet and orange), p = 0.29; data vs. N_V1_/N_RGC_ = 2.5 (green and orange), *p < 0.01; two-sample Kolmogorov-Smirnov test). Thus, our model also provides a rationale for the observed topographic correlation of salt-and-pepper organization in rodents.

Next, we examined the exact condition of the mapping ratio that generates columnar and salt-and-pepper organizations, respectively. For this, we applied a mathematical model for Nyquist sampling^18^, to investigate how topographic information of underlying retinal mosaics pattern could be differently mapped onto cortical space, depending on the mapping ratio condition. We first estimated a topographic cluster pattern of orientation tuning in the mosaics in Figure 2a^11^, from estimation of the local profile of ON and OFF RGC receptive fields (Fig. 2e, left). Then, the spatial periodicity of this retinal cluster was measured by FFT analysis of the filtered orientation tuning map (Period = 431 µm; see Extended Data Fig. 3 for details). Using the estimated period of the tuning map in the retinal source pattern, we defined the sampling ratio, *Γ*, as the number of V1 neurons (n_V1_) in a single spatial period. According to the Nyquist theorem^18^, the sampling condition *Γ* < 2 will cause an aliasing problem, thus our model predicts that parametric division of cortical organizations will occur around the threshold *Γ* = 2. A high retina-to-cortex sampling ratio *Γ* (large V1) would develop a periodic orientation map by projecting RGC periodicity into the cortex with no sampling alias (Fig. 2e, right, *Γ* = 6), while low sampling ratio *Γ* (small V1) would cause a noticeable aliasing of RGC periodicity. This produced more irregular clustering of orientation tunings in neighbouring V1 neurons (Fig. 2e, right, *Γ* = 1.5).

To prove that our observation of the parametric division between columnar and salt-and-pepper organization in animal data was mathematically predicted by the Nyquist theorem, we performed model simulations on the cortical organization for various sampling ratios. As predicted, local map continuity (or degree of spatial clustering) of orientation tuning increased with increase in the retina-to-cortex sampling ratio Γ (Fig. 2f), indicating the transition from salt-and-pepper organization to a smooth columnar map. Importantly, we observed that this transition appeared to change abruptly around the sampling ratio *Γ* = 2, the Nyquist frequency. This result explains why the cortical organizations observed so far are either columnar or salt-and-pepper, but without intermediates between these two stages.

Finally, we estimated the sampling ratio Γ for diverse mammalian species from the ratio between the number of neurons in V1 and in the retina, and compared it with the V1 organization of orientation tuning in each species (Fig. 2f and 2g). As predicted, the group of species with or without columnar clustering was successfully distinguished by the sampling ratio predicted from the Nyquist theorem.

## Discussion

Our findings suggest that both columnar and salt-and-pepper organizations of cortical circuits originate from retinal mosaics and by a universal development process. A mathematical sampling model shows that retino-cortical mapping is a prime determinant of the topography of cortical organizations and this prediction was confirmed by neural parameter analysis of data from eight species. This result implies that evolutionary variation of the size of retina and V1 has led to development of distinct topographical circuitry in V1, without species-specific developmental principles.

One might argue that the development of salt-and-pepper organization in rodents might originate from other characteristics of the rodent visual system. For example, it was reported that mice have diverse types of RGCs (> 30 types)^19^. However, the majority of these types of RGC is still classified as ON or OFF classes of RGC and each single class of RGC has been observed to be tiled regularly across the retinal surface^20^ as in higher mammals, implying that the regularly structured RGC mosaics may develop cortical organizations in both higher mammals and rodents, resulting in blueprints for cortical topography.

The large convergence in the retino-thalamic pathway in rodents is another factor to be considered. Electron microscopy techniques revealed that axons from several dozen RGCs anatomically innervated an LGN neuron in a mouse^21^, while most LGN neurons in cats appear to receive their major input from only 1–2 RGCs^22^. Therefore, this greater convergence in rodents has been considered to interrupt the development of a columnar organization in V1^14,23^. However, in a recent study on mice in which the functional connectome between retina and LGN was examined, it was reported that the response of LGN neurons is also mainly modulated by two major types of RGCs^24^ and that this is similar in primates. In addition, the receptive fields of mouse V1 neurons mostly consist of a couple of ON and OFF subregions^25^, implying that the functional connectome of retino-cortical pathways might be more similarly structured across species than is predictable from an anatomical connectome.

Previous studies also suggest that the emergence of orientation tuning in the earlier stage of visual pathway in rodents might provide the source of the salt-and-pepper organization^26^. However, the major thalamic inputs to V1 still consist of untuned units^27^, thus the LGN orientation-selective cells do not provide the primary source of orientation tuning in V1. In addition, the receptive field of most orientation-selective V1 neurons consist of 1-2 ON and OFF sub-regions^28^ as in primates. Considering that feedforward inputs for thalamic neurons are from retinal afferents, all these results suggest that the spatially biased arrangement of ON and OFF retinal inputs might be the major source of the orientation tuning, both in rodents and higher mammals.

The functional role of the columnar and salt-and-pepper organization has been debated^29,30^ and future studies are needed to reveal whether each type of organization is the optimized form of functional circuits under different physical constraints in each species. Our findings may provide advanced insight into the study of distinct cortical architectures under a universal principle of development process.

## Methods

### Connectivity and receptive field models

To model a V1 receptive field, we assumed that the RGCs are statistically wired to local V1 neurons around the corresponding cortical location^31^. The connection weight (*w*) of the retino-cortical projection was constrained by a 2D Gaussian distribution with a standard deviation (*σ*_*con*_) of 23 µm, so that a couple of ON and OFF RGCs strongly contributed to form a receptive field of a V1 neuron^31^.

We assumed that parameters for the retino-cortical projection could be approximated as quasi-consistent in the local eccentric area. In particular, we assumed that every cortical cell receives input from a similar number for RGCs regardless of variation in RGCs density and retino-cortical magnification, because the density of the RGCs and retino-cortical magnification are proportional across retinal eccentricity in the observed RGC data at large scale^32,33^ (Extended Data Fig. 1). Synaptic weighting between *i*^*th*^ RGC and *j*^*th*^ cortical sites (*w*_*ij*_) was defined as

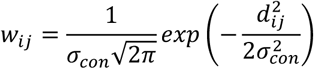

where *d*_*ij*_ represents the distance from the centre of the *i*^*th*^ RGC to the projected location of the *j*^*th*^ cortical site.

The receptive fields of RGCs were defined as a centre-surround 2D Gaussian model. The standard deviation of the surrounding region was set to three times that of the centre region^34^. The receptive field of the *i*^*th*^ RGC (*Ψ*_*i,RGC*_) was defined as

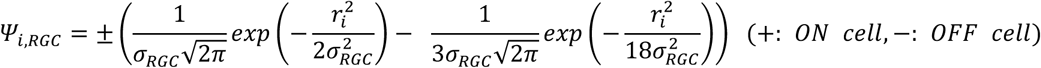

where *r*_*i*_ is a distance vector from the centre of the *i*^*th*^ RGC to each position of the visual field and *σ*_*RGC*_ was set as 43.1 µm for ON and 38.4 µm for OFF RGC. The receptive fields of V1 neurons were defined by the linear sum of the receptive fields of connected RGCs, so the receptive field of *j*^*th*^ cortical site (*Ψ*_*j,v*1_) was defined as

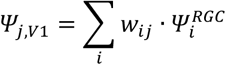

### Measurement of cortical functional tuning

The preferred orientation and selectivity of the calculated V1 receptive field was estimated from its Fourier transform ***Ψ***(***ω***). The preferred orientation (***θ***_***pref***_) and preferred spatial frequency (***ω***_***pref***_) were defined as ***θ***_***pref***_ = **arg**(***μ***) /2 and ***ω***_***pref***_ = |***μ*** |, where

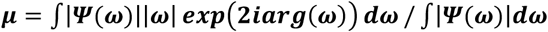

The orientation selectivity index (OSI) was defined as

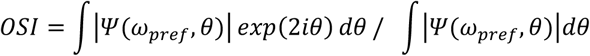

The tuning similarity between two cortical neurons was obtained by measuring a Pearson correlation coefficient of their receptive field in the frequency domain. The tuning similarity was measured from all the pairs of cortical neurons and was averaged over distance. Receptive fields were Fourier transformed and the non-linear response of neurons, *R*_*f*_, was estimated by applying a sigmoidal kernel,

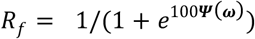

### Experimental data

The anatomical data used in this study were from ref. 35–38 for macaque, from ref. 8,37–40 for cat, from ref. 8,38,41– 43 for ferret, from ref. 38,44,45 for tree shrew, from ref. 37,38,46,47 for rabbit, from ref. 37,38,48,49 for mouse, from ref. 8,37,50,51 for rat. The number of V1 neurons in each species was estimated from the cell density (average cell distance, *d*_*v*1_ = 25 µm)^52^ and the size of V1.

## Acknowledgments

This work was supported by the National Research Foundation of Korea (NRF) grant funded by the Korea government (MSIT) (No. NRF-2016R1C1B2016039, NRF-2019R1A2C4069863, NRF-2019M3E5D2A01058328) (to S.P.).

## Author contributions

S.P. conceived the project. J.J., M.S. and S.P. designed the model. J.J., M.S. performed the simulations. J.J., M.S. and S.P. analysed the data. J.J., M.S. and S.P. wrote the manuscript.

---

**Extended Data Table 1.**
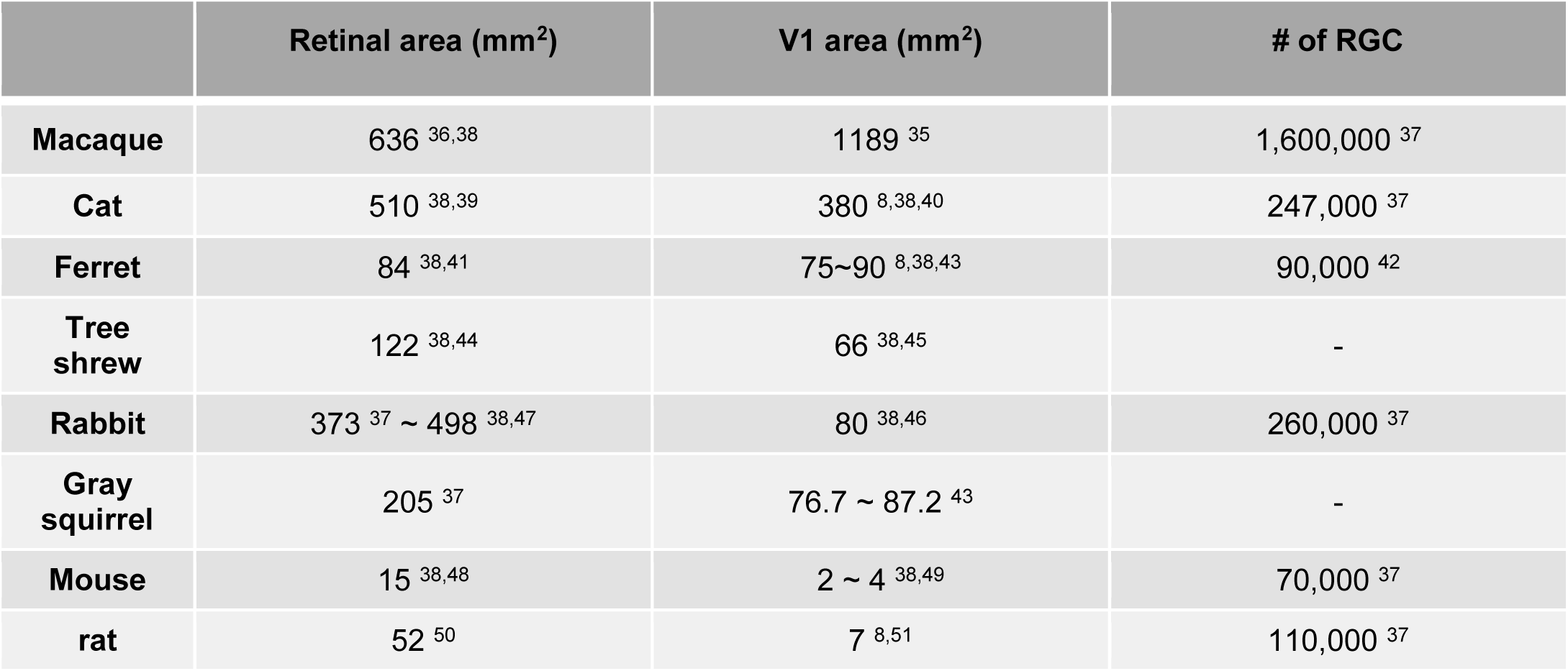
Experimental data of retina and V1 anatomy in diverse species.

**Extended Data Fig.1.**
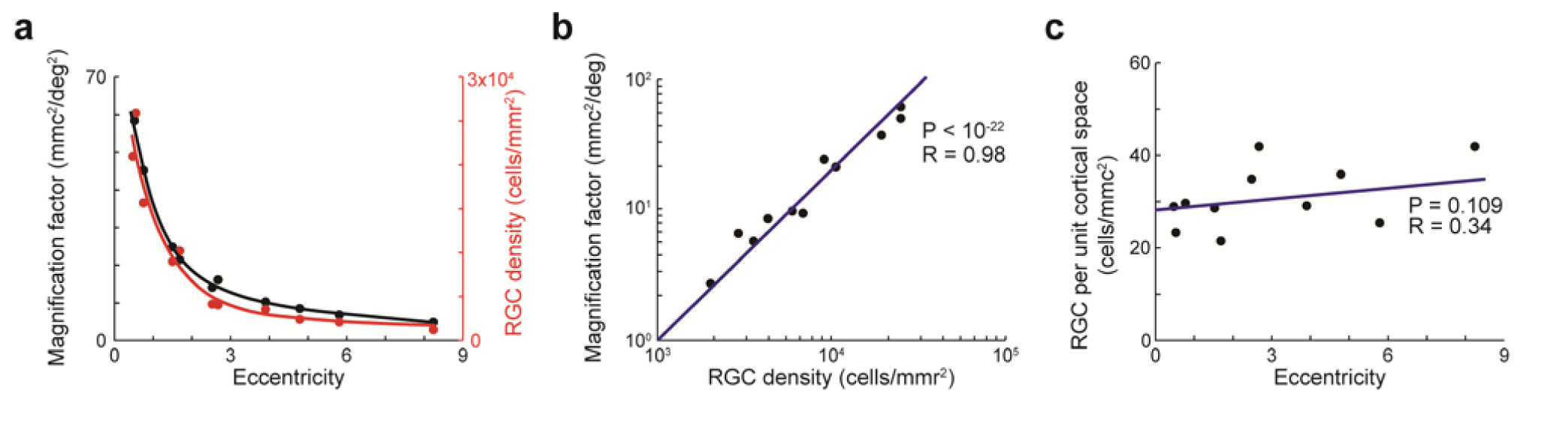
Retino-cortical mapping approximated as quasi-consistent across retinal eccentricity. **a**, Both the density of the RGCs and retino-cortical magnification factor change together as a function of eccentricity^32^ (the large magnification in the dense RGC areas of the retina, and small magnification in the sparse areas). **b**, RGCs density and magnification factor are directly proportional for each eccentricity in (a), indicating strong linear correlation. **c**, As a result, the number of RGC per unit cortical space could be approximated as quasi-consistent, implying that every cortical cell receives input from a similar number of RGCs regardless of variation in cell density^33^. In the graphs, mmr and mmc indicate mm in retinal and cortical space, respectively.

**Extended Data Fig.2.**
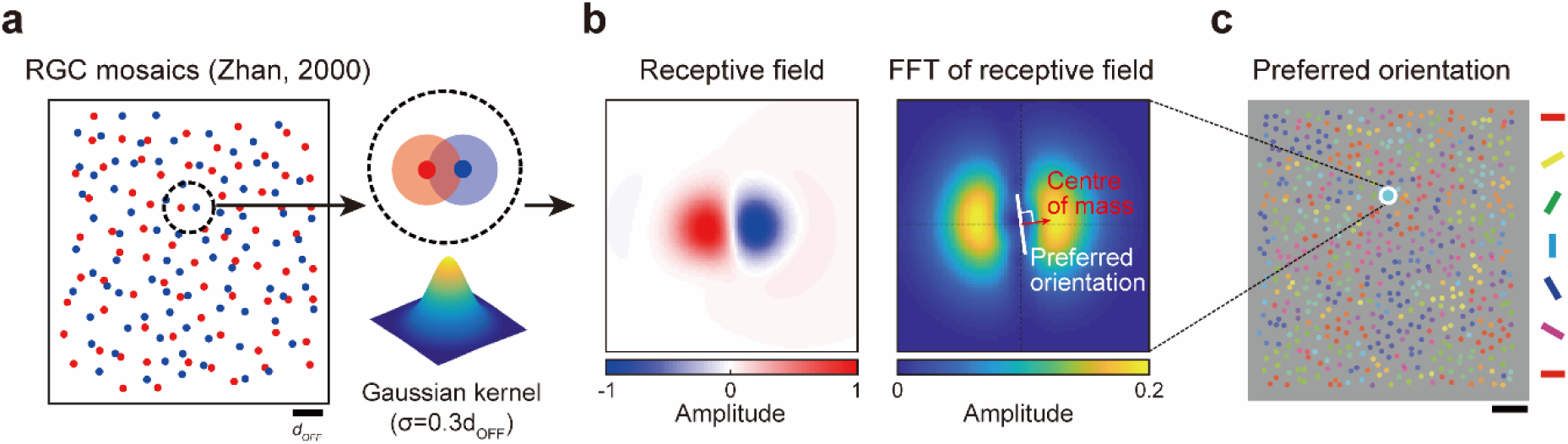
Estimation of orientation tuning in local V1 area from retinal mosaics. **a**, Cortical pooling (connection probability and strength) of retinal ganglion cells (RGCs) by a V1 neuron was modelled as a 2-D Gaussian kernel^12,31^. **b**, *Left*, Structure of local V1 receptive fields was simulated as a weighted sum of ON and OFF RGC receptive fields. *Right*, Orientation tuning of the V1 receptive field was determined using FFT analysis. The preferred orientation was found to be an orthogonal angle of the vector between the origin and the centre of mass. **c**, Orientation tuning map estimated from local orientation tuning at each cortical site.

**Extended Data Fig.3.**
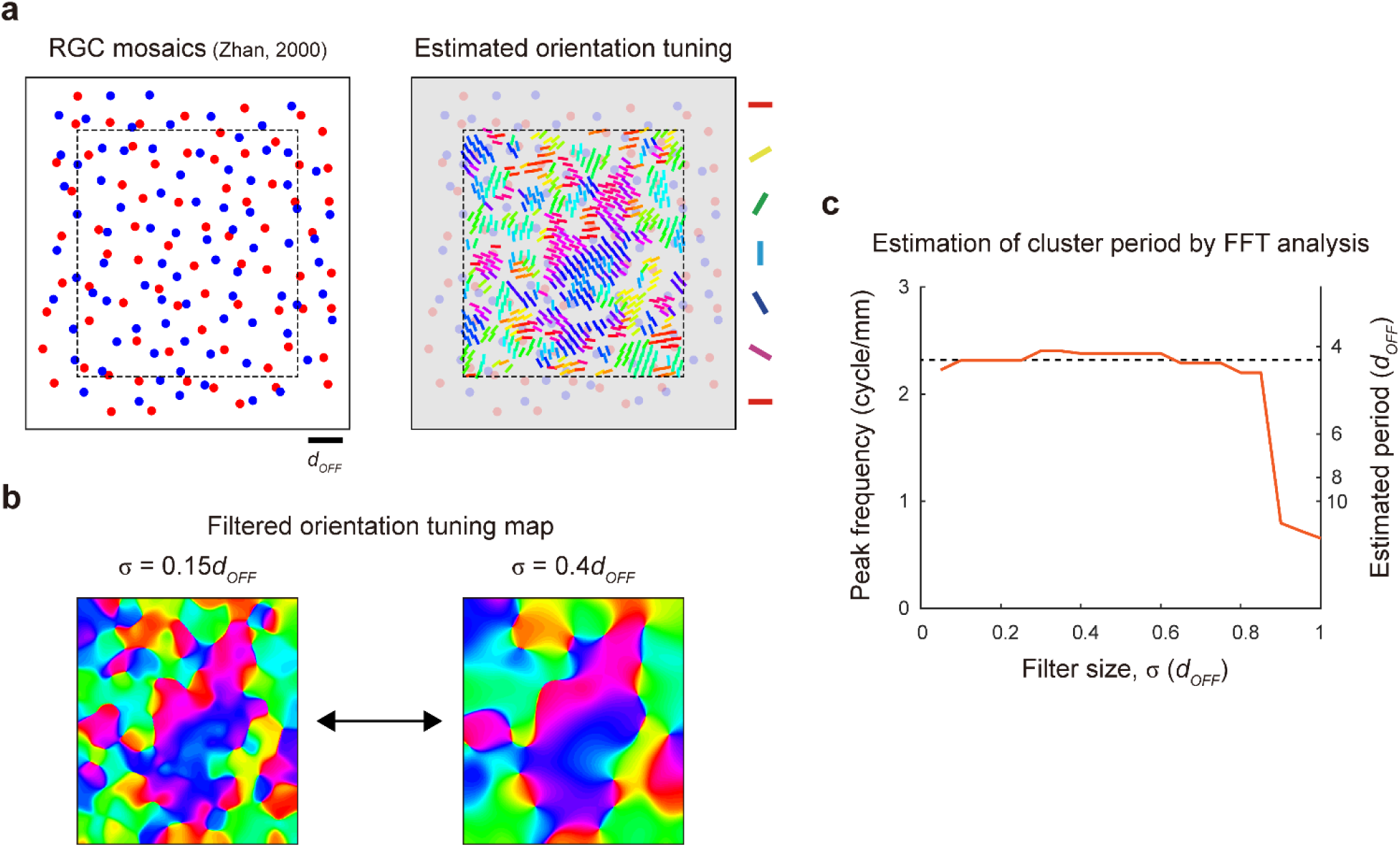
Robustness of spatial periodicity in retinal mosaics. **a**, Orientation tuning induced from the local arrangement of ON and OFF RGCs was estimated^53^. Boundary region (width = 1 *d*_*OFF*_) was excluded for more accurate estimation of local orientation tuning. **b**, Estimated organization of the orientation maps smoothed by 2D Gaussian filters of different sizes. **c**, To find a consistent spatial period of the map, the peak frequency calculated from FFT analysis of maps was measured for various filter sizes. Black dashed line indicates the average of the peak frequency within a plateau (2.32 cycles per mm). Thus, the spatial period observed in the retinal mosaics is 1 / 2.32 (cycle/mm) = 0.431 mm = 4.23 *d*_*OFF*_ (*d*_*OFF*_ = 0.102 mm).

